# Natural alleles at the *Doa* locus underpin evolutionary changes in *Drosophila* lifespan and fecundity

**DOI:** 10.1101/2022.10.14.512079

**Authors:** Katja M. Hoedjes, Hristina Kostic, Laurent Keller, Thomas Flatt

## Abstract

‘Evolve and resequence’ (E&R) studies in *Drosophila melanogaster* have identified many candidate loci underlying the evolution of ageing and life history, but experiments that validate the effects of such candidates remain rare. In a recent E&R study we have identified several alleles of the LAMMER kinase *Darkener of apricot* (*Doa*) as candidates for evolutionary changes in lifespan and fecundity. Here, we use two complementary approaches to confirm a functional role of *Doa* in life-history evolution. First, we used transgenic RNAi to study the effects of *Doa* at the whole-gene level. Ubiquitous silencing of expression in adult flies reduced both lifespan and fecundity, indicating pleiotropic effects. Second, to characterize segregating variation at *Doa*, we examined four candidate single nucleotide polymorphisms (SNPs; *Doa-1*, −*2*, −*3*, −*4*) using a genetic association approach. Three candidate SNPs had effects that were qualitatively consistent with expectations based on our E&R study: *Doa-2* pleiotropically affected both lifespan and late-life fecundity; *Doa-1* affected lifespan (but not fecundity); and *Doa-4* affected late-life fecundity (but not lifespan). Finally, the last candidate allele (*Doa-3*) also affected lifespan, but in the opposite direction than predicted.

## 1. Background

‘Evolve & resequence’ (E&R) studies, combining experimental evolution experiments with whole-genome sequencing, have emerged as a powerful method for identifying the genetic basis of evolutionary change [1–5]. In the *Drosophila melanogaster* model, for example, E&R studies have been successfully used to identify candidate loci underlying thermal adaptation [6, 7], the evolution of developmental rate [8], body size [9], hypoxia tolerance [10, 11], courtship song [12], lifespan and late-life fertility [13–15], dietary metabolism [16], pathogen resistance [17, 18], egg size [19], desiccation resistance [20], starvation resistance [21], and factorial selection on multiple life-history traits [22], among others.

Despite the identification of many putatively adaptive loci in such E&R studies, experimental assays that validate the presumed functional effects of such candidates are rare, which remains a major challenge for current tests of adaptation at the genetic level [1–3, 23]. Here, we examine the putative life-history effects of a candidate locus, *Darkener of apricot* (*Doa*), and associated candidate alleles which we have previously identified in an E&R experiment on longevity and late-life fertility in *D. melanogaster* [22]. *Doa*, a member of the LAMMER kinases, is known to phosphorylate a wide range of substrates and to be involved in many biological functions, such as embryonic development, oocyte formation, somatic sex determination, courtship behavior, and oxidative stress resistance (see Supplementary file 1 for additional information) [24–30]. Interestingly, *Doa* has been identified as a promising life-history candidate locus in several independent E&R experiments and genome-wide association studies (GWAS) on lifespan and late-life fertility, egg volume, and ovariole number [15, 19, 22, 31–33]. These findings make *Doa* a prime candidate locus for further study. Importantly, while several gene functions of *Doa* have been well established in molecular genetics studies (see above), the putative effects upon fitness components of naturally occurring polymorphisms at *Doa* have not yet been assessed.

In our experiments, we used two complementary approaches to investigate the functional role of *Doa* in affecting two major fitness components that had evolved in our E&R study, lifespan and fecundity [22, 34]. First, we used ubiquitous adult-specific RNAi silencing to examine the overall effects of *Doa* on lifespan and fecundity. Similar to using null mutants (amorphic mutations), this type of functional test is aimed at understanding function at the level of the whole gene; it can potentially reveal the complete phenotypic effects of a candidate gene, including any pleiotropic (and potentially deleterious) functions that it might have (e.g., [35, 36]). Given that specific alleles or mutations of a pleiotropic gene may differ in the functions they affect, however [35–37], we next investigated segregating variation at *Doa* by examining four candidate SNPs using a genetic association approach based on ‘Mendelian randomization’. Beyond confirming a role of the *Doa* locus in life-history adaptation, our results illustrate key differences in the effects of the four investigated SNPs on fecundity and longevity.

## 2. Material and methods

### (a) Transgenic RNAi

Transgenic *in vivo* RNAi was performed using the mifepristone (RU486)-inducible GeneSwitch(GS)-GAL4 system in combination with *Doa* UAS-RNAi constructs [38]. The GS system allows us to drive the expression of *Doa* UAS-RNAi during the adult stage only, thereby avoid potential developmental carry-over effects. Importantly, this system makes it possible to compare the effects of RNAi (i.e., the application of the drug, resulting in RNAi-mediated knockdown) to a negative control (i.e., no application of the drug) within the same transgenic genotype, thus providing a maximally robust control with regard to genotype possible. We used the ubiquitously expressing *daughterless* (*da)*-GeneSwitch(GS)-GAL4 construct [39] (courtesy of Véronique Monnier, Paris) to drive the expression of two independent *Doa* UAS-RNAi lines, obtained from the Vienna *Drosophila* RNAi Center (VDRC) (#19066 [D19]; #102520 [D10]), with construct insertions on chromosome 3 and 2, respectively, thus controlling for potentially confounding effects of insertion position. These constructs target the catalytic domain of *Doa* that is shared by all isoforms.

All lines were kept and assays were performed at 25°C, 65% humidity and 12h:12h light:dark cycle, on a cornmeal-yeast-sucrose-agar medium (per 1 liter of food: 7 g agar, 50 g sucrose, 50 g cornmeal, 50 g yeast, 6 ml propionic acid, 10 ml of a 20% nipagin stock solution). After emergence, flies were kept on medium containing either 100 or 200 μg/ml mifepristone (233 or 466 μM, respectively) dissolved in ethanol, or on control medium (i.e., containing ethanol without mifepristone). Note that increasing levels of mifepristone can induce higher levels of gene knockdown, in a dose-dependent fashion. We confirmed that both RNAi lines resulted in a significant knockdown of *Doa* expression using quantitative real-time PCR (qRT-PCR) (see Supplementary File 2 for details).

Mifepristone concentrations of up to 200 μg/ml have previously been used without detrimental effects on survival of adult flies [15, 39, 40]. To confirm this, we tested the effect of 100 and 200 μg/ml mifepristone on lifespan and fecundity of F1 flies of a cross of *da*-GS-GAL4 females with males of the isogenic progenitor strain for the VDRC (GD) RNAi library strains, *w*^1118^ (#60000). We did not observe any confounding deleterious effects of these concentrations on the phenotypes of interest (Supplementary File 3).

To assess the effect of RNAi directed against on *Doa* on lifespan, cohorts of F1 offspring between crosses of *da*-GS-GAL4 virgin females and males carrying one of the two UAS-RNAi constructs or the isogenic control strain were collected within a 24-hour window. Flies were sexed under mild CO_2_ exposure and transferred to 1-liter demography cages with food vials (with 0, 100, or 200 μg/ml mifepristone) attached to the cages. For each genotype and mifepristone concentration, we set up three replicate cages, each containing 75 flies per sex. Dead flies were scored and fresh food was provided every two days. Differences in lifespan between mifepristone-induced RNAi and uninduced controls were analyzed in *R* (v.3.3.1) using mixed-effects Cox (proportional hazards) regression with mifepristone concentration, sex and their interaction as fixed effects and with ‘replicate cage’ as a random effect, using the *R* package *coxme* (v2.2-5).

The effect of *Doa* RNAi on fecundity was assessed by measuring daily egg production of females. Virgin females were collected from the F1 offspring of *da*-GS-GAL4 females and males carrying one of the two UAS-RNAi constructs or the isogenic control strain. After 24 hours, two virgin females and two *w^1118^* males were placed together in vials with either 0 (i.e., control) or 200 μg/ml mifepristone. Ten replicate vials were prepared per genotype and mifepristone concentration. Flies were left for 48 hours to ensure mating and consumption of mifepristone before the start of the experiment; after this period, they were transferred to fresh vials with food (with 0 or 200 μg/ml mifepristone, respectively) to lay eggs for 24 hours. The number of eggs laid by each pair of females was counted under a dissecting microscope; daily egg production was measured for nine consecutive days. We calculated average fecundity per female over 3 days in order to average out day-to-day variation in egg laying. Fecundity data were analyzed in *R* (v.3.3.1) using generalized linear mixed models with a Poisson distribution and with mifepristone concentration, day and their interaction as fixed effects and with ‘replicate vial’ as a random effect using the *R* package *lme4* (v.1.1-13). Exposure to mifepristone of the control line did not cause adverse (and thus confounding) effects on lifespan and fecundity (see Supplementary File 3)

### (b) SNP association study

To examine whether the four experimentally candidate SNPs at *Doa* affect lifespan and fecundity (see below for details of SNP identification), we performed a genetic (SNP) association study, based on ‘Mendelian randomization’ [41–43]. Mendelian randomization (MR) approaches aim to identify putative causal effects of candidate loci by testing the alternative allelic states in a genetically diverse background to limit the impact of potentially confounding (epistatic) factors in the genetic background. A critical factor for the reliability of MR approaches is the lack of linkage disequilibrium (LD) between the focal locus and other loci in the genetic background. Here, we used strains from the *Drosophila* Genetic Reference Panel (DGRP [44]; obtained from the Bloomington *Drosophila* Stock Center [BDSC]), which provides ample natural genetic variation for MR. As linkage disequilibrium (LD) typically decays very rapidly in *D. melanogaster*, within a few hundred base pairs or so [44], the MR approach is expected to provide information on the functional impact of an individual candidate SNP, with little or no confounding effects of other (*Doa*) SNPs. To confirm this, we analyzed LD (measured by pairwise *r*^2^) among all polymorphic *Doa* SNPs (minor allele frequency ≥ 0.1) in the complete panel of DGRP lines, as done by [43] before (see Supplementary File 4).

For each of the four candidate nucleotide positions at *Doa*, we randomly selected 20 distinct DGRP lines that were fixed for the SNP allele which was previously identified as being the major allele in the short-lived, early-reproduction experimental evolution lines (control lines; see [22]). Similarly, we randomly selected 20 lines fixed for the SNP allele that was found to be the major allele in the long-lived, late-reproduction experimental evolution lines (see Supplementary Table 3 for details of the crosses). SNP genotypes were confirmed by sequencing a small genomic region surrounding each SNP using Sanger sequencing. For each allelic state (‘short-lived’ vs. ‘long-lived’) and nucleotide position, we generated 10 unique F1 crosses, each cross being made from a different pair of distinct DGRP lines sharing the same SNP state (i.e., virgin females of one strain crossed to males from the other strain); because the DGRP lines are inbred, this was done to minimize potentially confounding homozygous effects at non-candidate loci in the genomic background. Thus, for each of the four candidate nucleotide positions and for each alternative allele (‘short-lived’ vs. ‘long-lived’) we had a 10-fold replicated panel of independent F1 genotypes fixed for a given SNP allele but maximally heterozygous at other genomic positions. For details see the overview in Supplementary Table 3; in the end, due to the low viability of some DGRP lines and F1 crosses, we phenotyped between 8-10 F1 crosses per candidate SNP and allelic state. To evaluate whether any other positions in the genome, besides the candidate SNP allele, were highly differentiated between the four pairs of panels, we calculated SNP-wise F_ST_ based on the method of Weir and Cockerham [45], using the pooled genome sequence information per panel [42] (Supplementary File 5).

F1 crosses were reared and assays performed at 25°C, 65% humidity and 12h:12h light:dark cycle, on a cornmeal-yeast-sucrose-agar medium, as described above. Lifespan was measured using demography cages, as above. Flies that had emerged within a 24-hour window were collected, and for each F1 cross 75 males and 75 females were placed in a single demography cage. Differences in lifespan between the two allelic states of each SNP were analyzed in *R* (v.3.3.1) using mixed-effects Cox (proportional hazards) regression with allele, sex and their interaction as fixed effects and with ‘F1 cross’ as a random effect using the *R* package *coxme* (v2.2-5).

Fecundity was measured over a period of 30 days after eclosion in order to provide insight into early (peak) and late (post-peak) fecundity. Flies that had eclosed within a 24-hour window were collected for crosses; for each F1 cross, two females and two males were placed in a vial containing regular medium, with three replicate vials per F1 cross. Every third day (i.e., on days 3, 6, 9, 12, 15, 18, 21, 24, 27 and 30), the number of eggs laid by each pair of females during a 24-hour period was determined using a dissecting microscope. Fecundity was analyzed using generalized linear mixed models with a Poisson distribution in *R* (v.3.3.1), with allelic state, day and their interaction as fixed effects and ‘replicate vial’ as a random effect using the *R* package *lme4* (v.1.1-13).

## 3. Results and Discussion

### (a) Evolutionary changes at *Doa* might underpin life-history evolution

We previously identified *Doa* as a life-history candidate locus in an E&R experiment in which fruit flies were selected for late-life fecundity and where a longer lifespan evolved as a correlated response [22]. In that study, *Doa* was identified as one of multiple candidate loci under selection (Figure 1A). Genetic tests of candidate genes and SNPs are critical to distinguish the putative causal loci from false positive signals and to determine if they have an effect on either one or both of the evolved traits. *Doa* is a very large gene, spanning 34.7 kb, and encodes at least 6 protein isoforms that are expressed in an age- and tissue-specific manner from alternative promotors and which have different, non-redundant functions [25, 26]. SNPs located at different positions within the *Doa* gene might thus have different functions.

**Figure 1.**
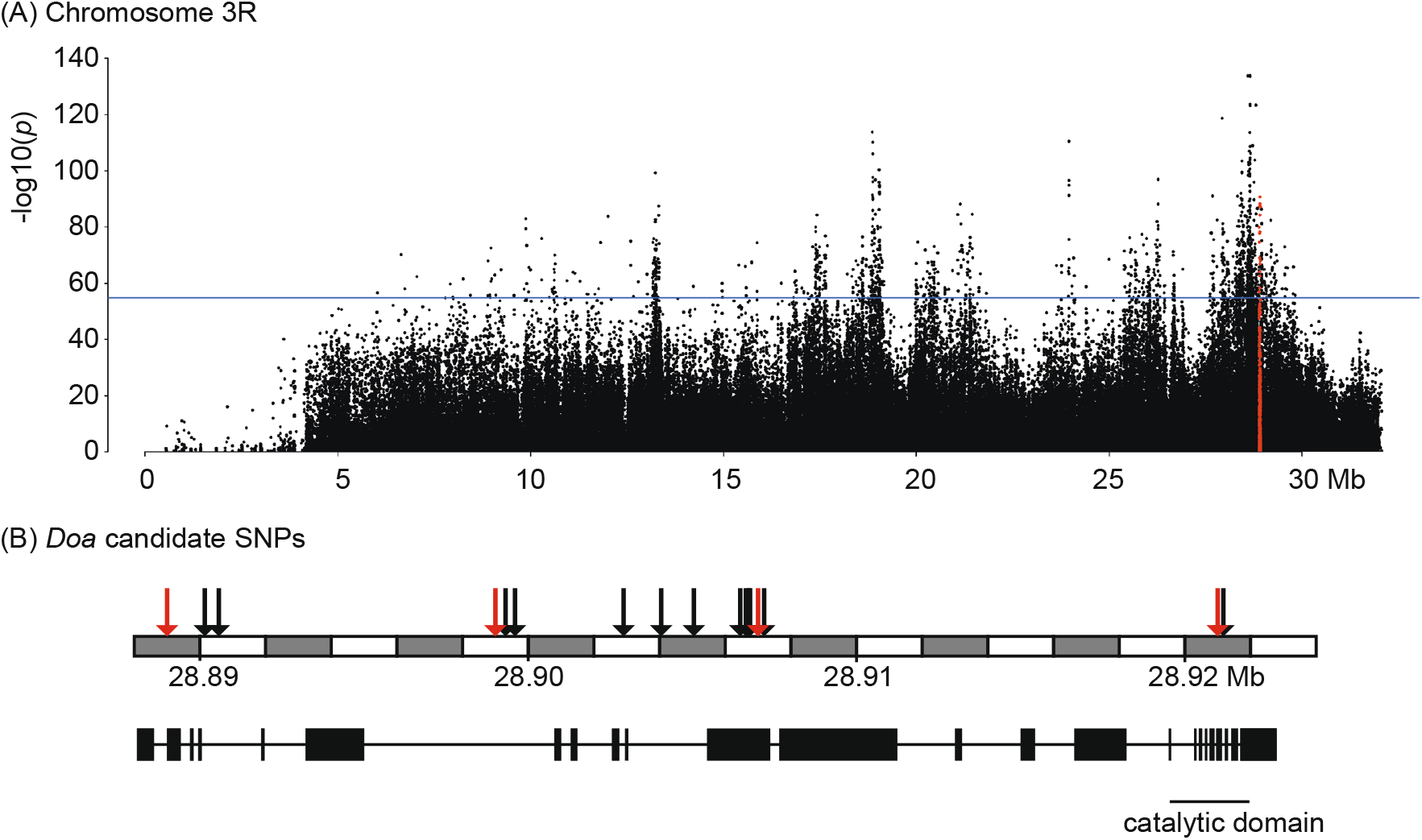
Overview of the experimentally evolved *Doa* SNPs. (A) The Manhattan plot demonstrates the position of *Doa* on chromosome 3R in the Evolve-and-Resequence study by Hoedjes *et al*. [22]. *Doa* (all SNPs are highlighted with red) is located within a relatively large region at the end of chromosome 3 that had been under selection. The horizontal blue line indicates the threshold for significant candidate SNPs (FDR=0.0005) (B) Shows the gene structure of *Doa* with the 16 significant candidate SNPs (at FDR<0.0005; indicated with arrows) that were identified as candidates for ageing and reproduction. Red arrows indicate the four SNPs that have been investigated functionally in this study. Coordinates according to reference genome v6.

**Figure 2.**
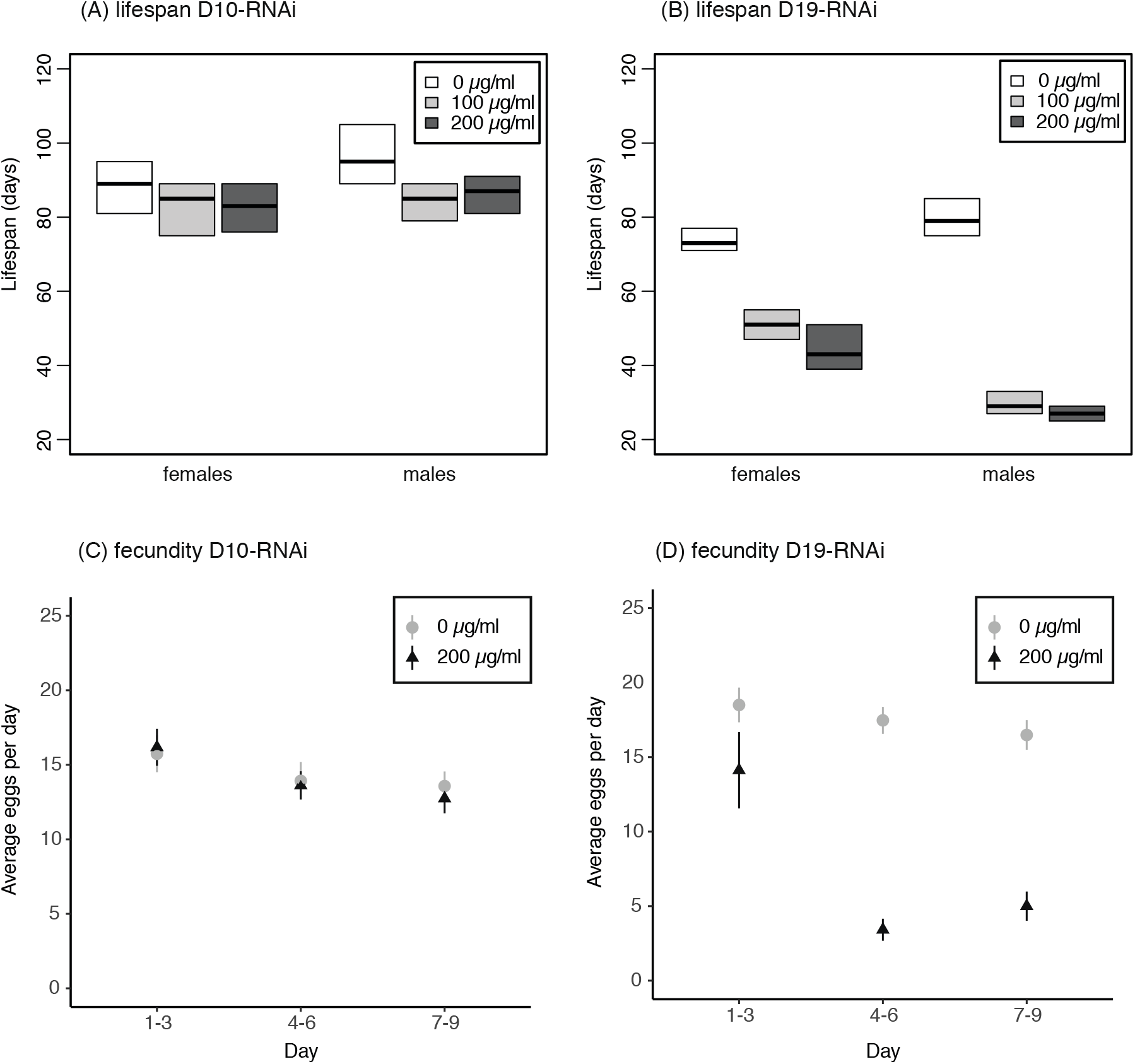
Knockdown of *Doa* using transgenic RNAi demonstrates significant effects on lifespan and fecundity. Two independent RNAi constructs (D19 and D10) that both target the catalytic domain of *Doa* were used in combination with the mifepristone-inducible GeneSwitch-Gal4>UAS system. Concentrations refer to the concentrations of mifepristone (RU-486) used to induce RNAi. A significant reduction of adult lifespan was observed with both constructs (A: construct D10, B: construct D19), whereas a reduction in female fecundity was observed for construct D19 only (C: construct D10, D: construct D19) (see Table 1 for statistics).

In total, we identified 16 biallelic SNPs (either intronic or in the coding region) having highly significant allele frequency differentiation (FDR <0.0005) between the early-reproduction and late-reproduction (and hence increased lifespan) selection regimes in our E&R study [22] (Figure 1B, Supplementary Table 1). We chose four of these SNPs, based on their allele frequencies differentiation between the selection regimes and their distribution across the length of the gene, for functional assays (Figure 1, red arrows, and Table 2; see below). As these SNPs were identified in an E&R study performed in a controlled laboratory setting, we first compared their allele frequencies to those in natural populations, using data from the DEST database [46] (see Supplementary File 6 for approach). These analyses showed that all four SNPs are polymorphic in European populations and that the allele frequencies observed in the E&R study fall within the range of frequencies in natural populations (*Doa-1*: 0.16-0.57, *Doa-2*: 0.34-0.72, *Doa-3*: 0.65-0.98, *Doa-4*: 0.51-0.95, see Supplementary File 6 and Supplementary Table 2). Moreover, there was a significant latitudinal cline in allele frequency at *Doa-4* (χ^2^ = 174.3, *P_corrected_* = 0.0069) and a significant longitudinal cline at *Doa-3* (χ^2^ = 222.9, *P_corrected_* = 0.037) across European populations (Supplementary Table 4). This is notable because numerous life-history traits in *D. melanogaster*, including fecundity and lifespan, exhibit a clinal distribution, presumably due to spatially varying selection [3, 42, 43, 47–50]. These population genetic observations thus lend further support to the idea that *Doa* potentially represents a target of selection on life-history traits.

### (b) *Doa* has pleiotropic life-history effects

To functionally validate the role of *Doa* in longevity and fecundity at the level of the whole gene we knocked down all transcript variants by targeting the common, catalytic domain of *Doa* with ubiquitous (i.e., non-tissue-specific) transgenic RNAi. We employed two different RNAi constructs (i.e., two independent chromosomal insertions) to control for confounding effects of insertion position. To exclude potentially detrimental developmental carry-over effects of *Doa* knockdown on adult fitness, we reduced *Doa* expression levels in the adult stage only, by driving RNAi with the mifepristone-inducible GeneSwitch-Gal4 system [38]. For both *Doa*-RNAi constructs, we observed a significant reduction in lifespan in both sexes with increasing levels of mifepristone (Figures 1A-B, Table 1, Supplementary File 2). Overall, the effect on lifespan was strongest for the D19 construct, which also achieved a stronger knockdown of *Doa* as determined by qRT-PCR as compared to construct D10. For both constructs, there was a significant interaction between sex and mifepristone concentration, which reflects the overall stronger effects of *Doa* RNAi on male than female lifespan. These findings agree with the observation of Huang *et al*. [31] that weak, constitutive knockdown of *Doa* (also using construct D10, VDRC #102520) affects lifespan in males, but not females; however, in their study, the direction of the effect depended on assay temperature. Sexual dimorphism in longevity is not uncommon among organisms, inlcuding fruit flies [51, 52]. In terms of fecundity, we observed a strong, significant reduction in egg-laying rate for construct D19 but not for D10 (Figures 1C-D, Table 1).

**Table 1.** Statistical tests of the effects of *Doa* RNAi on lifespan and fecundity (egg laying rate). Two independent RNAi constructs (D19 and D10) that both target the catalytic domain of *Doa* were used in combination with the mifepristone-inducible GeneSwitch-Gal4>UAS system. A strain with the same genetic background as the two constructs (*w^1118^*) was used a control for adverse effects of mifepristone application. Significant effects are indicated in bold.

|  | Longevity |  |  | Fecundity |  |
| --- | --- | --- | --- | --- | --- |
| | $\chi^2$ | *P* | | $\chi^2$ | *P* |
| ***w<sup>1118</sup>* (control)** |  |  | ***w<sup>1118</sup>* (control)** |  |  |
| Mifepristone | 8.64 | 0.071 | Mifepristone | 7.31 | 0.063 |
| Sex | 95.68 | **<2.2e-16** | Day | 50.83 | **2.4e-10** |
| Interaction | 3.63 | 0.163 | Interaction | 3.59 | 0.166 |
| ***D10*** |  |  | ***D10*** |  |  |
| Mifepristone | 51.53 | **1.7e-10** | Mifepristone | 1.64 | 0.6512 |
| Sex | 83.56 | **<2.2e-16** | Day | 34.12 | **7.0e-07** |
| Interaction | 37.29 | **8.0e-9** | Interaction | 1.61 | 0.446 |
| ***D19*** |  |  | ***D19*** |  |  |
| Mifepristone | 317.24 | **<2.2e-16** | Mifepristone | 320.54 | **<2.2e-16** |
| Sex | 383.21 | **<2.2e-16** | Day | 508.72 | **<2.2e-16** |
| Interaction | 284.46 | **<2.2e-16** | Interaction | 307.51 | **<2.2e-16** |

**Table 2.** Details on the *Doa* candidate SNPs. The genomic location of the four *Doa* SNPs that were investigated functionally is shown. For each SNP, a significant difference in allele frequencies was observed between lines selected for early age-at-reproduction (“Control”) versus lines selected for late-life (postponed) reproduction (which associated with an evolutionary increase in lifespan: “Long-lived”).

| Name | Location | Feature | Allele frequency |  | Control allele | "Long-lived" allele |
| --- | --- | --- | --- | --- | --- | --- |
|  |  |  | Control | Long lived |  |  |
| Doa-1 | 3R:28'888'916 | intronic | 0.18 | 0.52 | C | T |
| Doa-2 | 3R:28'898'997 | intronic | 0.43 | 0.80 | A | G |
| Doa-3 | 3R:28'907'179 | missense | 0.55 | 0.90 | A | G |
| Doa-4 | 3R:28'921'024 | synonymous | 0.40 | 0.78 | C | T |

These findings show that the *Doa* gene has pleiotropic effects on lifespan and reproduction [22] and demonstrate that modifying expression in the adult stage is sufficient to mediate these effects. Moreover, the magnitude and the direction of these effects depend on the strength of the knockdown (also see [31]). To obtain a better understanding of the role of *Doa* in evolving populations we next studied the effects of the four candidate SNPs identified in the E&R study on fecundity and longevity (see above and [22]).

### (c) *Doa* natural alleles have pleiotropic and non-pleiotropic effects

The four candidate SNPs that were functionally characterized were located both in coding and non-coding regions; *Doa-4* is a synonymous SNP located in the catalytic domain that is shared by all isoforms; *Doa-1* and *Doa-2* are intronic SNPs, and may have regulatory functions; and *Doa-3* is a missense SNP located in exon N8 (N-terminal variable region), which encodes part of the 227 kD protein isoform [26]. To study the effects of these SNPs on fitness components we used a SNP association approach based on Mendelian randomization using lines of the *Drosophila Genetic Reference Panel* (DGRP) (see above, 2. Materials and methods).

For three of the *Doa* SNPs we found a significant correlation between allelic state and median lifespan (Figure 3, Table 3). For *Doa-1* and *Doa-2*, the correlation was in the predicted direction (i.e., increased median longevity of lines carrying the allele that was enriched in the long-lived E&R populations). Interestingly, there was also a significant correlation between lifespan and allelic state for *Doa-3*, but the direction was opposite to what we had hypothesized: although the “G” variant was the major allele in the long-lived experimental evolution lines [22], this allele was associated with lower median lifespan in our functional assays (Figure 3, Table 3).

**Figure 3.**
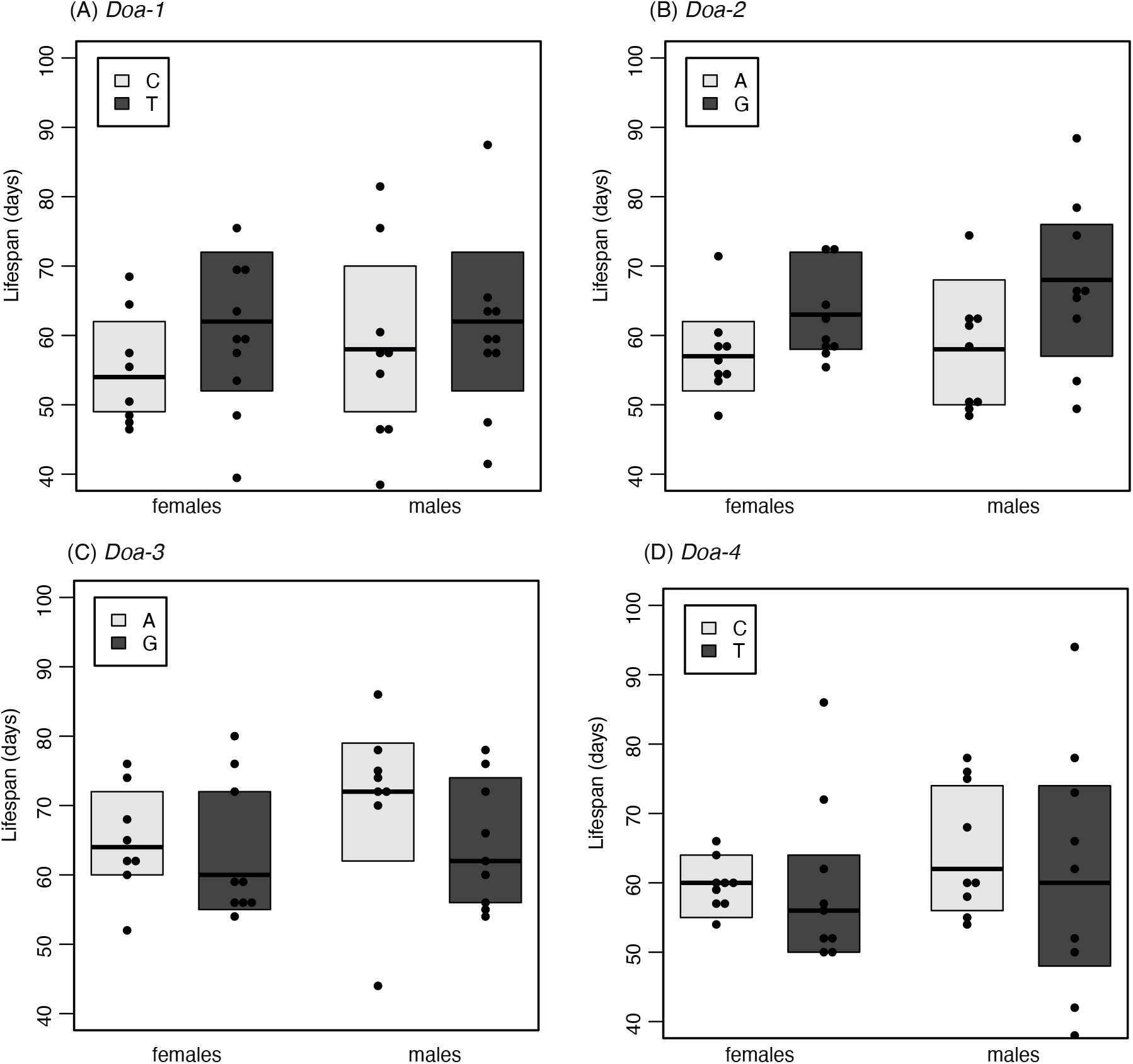
Association between candidate SNPs at *Doa* and lifespan. F1 crosses of the genetically diverse *Drosophila* Genetic Reference Panel with different allelic states were assessed to test the association of the candidate SNPs with lifespan. The graphs show the average lifespans of the F1 crosses for each allelic state at the four candidate *Doa* SNP positions investigated. The allele that is pre-dominantly found in long-lived, late-reproducing EE populations, as identified by Hoedjes et al. [22], is indicated by the darker shade. Significant differences in lifespan between the two allelic states were observed for *Doa-1*, *Doa-2*, and *Doa-3* (see Table 3 for statistics).

**Table 3.** Statistical tests of the effects of alternative alleles at the four *Doa* SNP positions on lifespan and fecundity (egg laying rate). F1 crosses of the genetically diverse *Drosophila* Genetic Reference Panel with different allelic states were set-up and tested for lifespan and fecundity in order to assess the association of the candidate *Doa* SNPs with lifespan.

|  | Lifespan |  |  | Fecundity |  |
| --- | --- | --- | --- | --- | --- |
| | $\chi^2$ | *P* | | $\chi^2$ | *P* |
| ***Doa-1*** |  |  | ***Doa-1*** |  |  |
| Allele | 43.27 | 4.02e-10 *** | Allele | 0.83 | 0.66 |
| Sex | 137.99 | <2.2e-16 *** | Day | 477.8 | <2.2e-16 *** |
| Allele*Sex | 41.42 | 1.23e-10 *** | Allele*Day | 0.24 | 0.6224 |
| ***Doa-2*** |  |  | ***Doa-2*** |  |  |
| Allele | 7.62 | 0.022 * | Allele | 16.23 | 0.0003 *** |
| Sex | 79.22 | <2.2e-16 *** | Day | 394.66 | <2.2e-16 *** |
| Allele*Sex | 3.35 | 0.067 | Allele*Day | 15.84 | 6.90e-05 *** |
| ***Doa-3*** |  |  | ***Doa-3*** |  |  |
| Allele | 12.02 | 0.0025 ** | Allele | 1.20 | 0.55 |
| Sex | 228.45 | <2.2e-16 *** | Day | 348.77 | <2.2e-16 *** |
| Allele*Sex | 11.46 | 0.00071 *** | Allele*Day | 1.13 | 0.28 |
| ***Doa-4*** |  |  | ***Doa-4*** |  |  |
| Allele | 3.78 | 0.15 | Allele | 9.42 | 0.009 * |
| Sex | 141.33 | <2.2e-16 *** | Day | 486.4 | <2.2e-16 *** |
| Allele*Sex | 3.17 | 0.075 | Allele*Day | 7.68 | 0.0056 * |

In terms of effects on fecundity, we observed a significant correlation between allelic state and egg-laying rates for two of the *Doa* SNPs, *Doa-2* and *Doa-4*. Moreover, in both cases, there was also a significant interaction between allelic state and age, indicating that the effect on fecundity was age-specific (Figure 4, Table 3). The difference in egg-laying rate became visible starting 18-24 days after eclosion, with a higher fecundity of the crosses carrying the alleles associated with selection for postponed reproduction in the E&R study [22].

**Figure 4:**
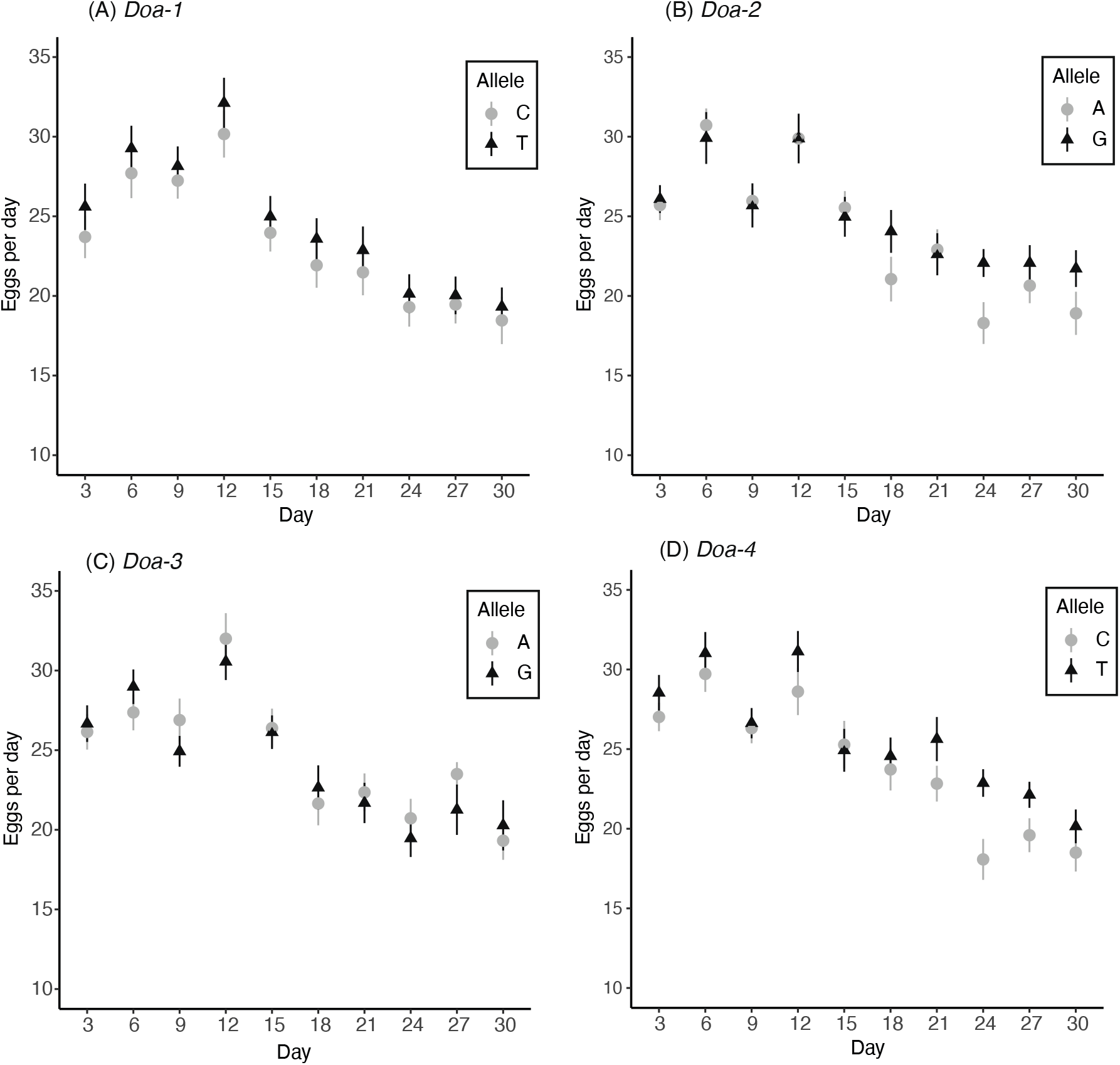
Association between candidate SNPs at *Doa* and fecundity. Shown are average egg laying rates over a period of 30 days after emergence of F1 crosses for each allelic state at the four *Doa* candidate SNP positions investigated. The allele that is pre-dominantly found in long-lived, late-reproducing EE populations, as identified by Hoedjes *et al*. [22], is indicated by the darker shade and triangles. Significant differences in lifespan between the two allelic states were observed for *Doa-2* and *Doa-4* (see Table 3 for statistics).

To rule out potentially confounding effects of linked causal SNPs in the genetic background, we analyzed LD (measured by pairwise *r*^2^) among *Doa* SNPs in the DGRP, which indicated very low levels of LD across the gene, as well as among the four candidate SNPs (see Supplementary File 4). In addition, analyses of genetic differentiation (SNP-wise F_ST_) among lines with alternative allelic states demonstrated that only the focal SNP was fixed (F_ST_ = 1) for each of the four panels of lines (Supplementary File 5). None of the other SNPs, both within *Doa* or elsewhere in the genome, were fixed between two panels of lines, and the number of strongly differentiated SNPs (F_ST_ > 0.5) was very low as well. These analyses strongly suggest that our findings are very unlikely to be confounded by LD or due to effects in the genetic background of the candidate SNPs tested.

## 4. Conclusions

Our study provides strong support for a role of *Doa* in the evolution of lifespan and fecundity in the fruit fly, as expected based on our previous E&R study [22]. Ubiquitous gene silencing of *Doa* using transgenic RNAi in adult flies reduced both lifespan and fecundity, indicating positive pleiotropy. In addition, each of the four candidate *Doa* SNPs tested had a significant effect on either lifespan and/or fecundity. The exact effects depended on the specific SNP, however, indicating that functional characterization of individual polymorphisms is essential for identifying the loci underlying adaptation. Three polymorphisms had effects on lifespan and/or fecundity that agree qualitatively well with predictions [22]. One of these SNPs, *Doa-2*, affected both lifespan and late-life fecundity, illustrating that even single nucleotide changes can have pleiotropic effects on complex traits (for other examples see [37, 42, 43]). However, one of the four SNPs, *Doa-3*, affected lifespan in the opposite direction than predicted. A possible explanation for this surprising result might be that this SNP, and potentially other SNPs at *Doa*, are part of functionally and evolutionarily important haplotypes subject to linkage disequilibrium and/or epistasis. Similar analyses of other candidate loci, both within *Doa* and elsewhere in the genome, could resolve these questions and provide a more comprehensive overview of the polygenic regulation of these traits. Together, our results illustrate that it is important to go beyond traditional gene knockdown or knockout analyses and to perform functional tests of putatively adaptive candidate loci in order to understand the genetic basis of evolutionary change.

## Data accessibility

The raw data for this paper is available at Dryad (https://doi.org/10.5061/dryad.0gb5mkm4t)

## Authors’ contributions

Definitions according to CRediT (https://casrai.org/credit/): <u>KMH</u>: Conceptualization, Data curation, Formal Analysis, Funding acquisition, Investigation, Methodology, Project administration, Supervision, Resources, Software, Validation, Visualization, Writing – original draft, Writing – review & editing; <u>HK</u>: Investigation, Writing – review & editing; <u>LK</u>: Conceptualization, Funding acquisition, Project administration, Supervision, Resources, Writing – review & editing; <u>TF</u>: Conceptualization, Supervision, Resources, Writing – review & editing.

## Acknowledgements

We are grateful to Tadeusz Kawecki (Lausanne) for generously letting the first author use his *Drosophila* facilities. We also thank Bas Zwaan (Wageningen) and the Flatt and Keller labs for discussion and support; Véronique Monnier for the da-GS-GAL4 strain; and the Bloomington Drosophila Stock Center (BDSC) and the Vienna Drosophila RNAi Center (VDRC) for providing stocks.

## References

1. Schlötterer, C., Kofler, R., Versace, E., Tobler, R., Franssen, S. U. 2014 Combining experimental evolution with next-generation sequencing: a powerful tool to study adaptation from standing genetic variation. Heredity (Edinb). 0, (10.1038/hdy.2014.86)

2. Schlötterer, C., Tobler, R., Kofler, R., Nolte, V. 2014 Sequencing pools of individuals - mining genome-wide polymorphism data without big funding. Nat Rev Genet. 15, 749–763. (10.1038/nrg3803)

3. Flatt, T. 2020 Life-History Evolution and the Genetics of Fitness Components in *Drosophila melanogaster*. Genetics. 214, 1–45. (10.1534/genetics.119.300160)

4. Long, A., Liti, G., Luptak, A., Tenaillon, O. 2015 Elucidating the molecular architecture of adaptation via evolve and resequence experiments. Nature Reviews Genetics. 16, 567–582. (10.1038/nrg3937)

5. Kessner, D., Novembre, J. 2015 Power Analysis of Artificial Selection Experiments Using Efficient Whole Genome Simulation of Quantitative Traits. Genetics. 199, 991–U147. (10.1534/genetics.115.175075)

6. Orozco-TerWengel, P., Kapun, M., Nolte, V., Kofler, R., Flatt, T., Schlötterer, C. 2012 Adaptation of *Drosophila* to a novel laboratory environment reveals temporally heterogeneous trajectories of selected alleles. Molecular Ecology. 21, 4931–4941. (DOI 10.1111/j.1365-294X.2012.05673.x)

7. Tobler, R., Franssen, S. U., Kofler, R., Orozco-Terwengel, P., Nolte, V., Hermisson, J., Schlötterer, C. 2014 Massive habitat-specific genomic response in *D. melanogaster* populations during experimental evolution in hot and cold environments. Mol Biol Evol. 31, 364–375. (10.1093/molbev/mst205)

8. Burke, M. K., Dunham, J. P., Shahrestani, P., Thornton, K. R., Rose, M. R., Long, A. D. 2010 Genome-wide analysis of a long-term evolution experiment with *Drosophila*. Nature. 467, 587–U111. (10.1038/nature09352)

9. Turner, T. L., Stewart, A. D., Fields, A. T., Rice, W. R., Tarone, A. M. 2011 Population-Based Resequencing of Experimentally Evolved Populations Reveals the Genetic Basis of Body Size Variation in *Drosophila melanogaster*. Plos Genet. 7, e1001336. (10.1371/journal.pgen.1001336)

10. Zhou, D., Udpa, N., Gersten, M., Visk, D. W., Bashir, A., Xue, J., Frazer, K. A., Posakony, J. W., Subramaniam, S., Bafna, V., et al. 2011 Experimental selection of hypoxia-tolerant *Drosophila melanogaster*. Proceedings of the National Academy of Sciences of the United States of America. 108, 2349–2354. (10.1073/pnas.1010643108)

11. Jha, A. R., Zhou, D., Brown, C. D., Kreitman, M., Haddad, G. G., White, K. P. 2016 Shared Genetic Signals of Hypoxia Adaptation in *Drosophila* and in High-Altitude Human Populations. Molecular Biology and Evolution. 33, 501–517. (10.1093/molbev/msv248)

12. Turner, T. L., Miller, P. M. 2012 Investigating Natural Variation in *Drosophila* Courtship Song by the Evolve and Resequence Approach. Genetics. 191, 633–U516. (10.1534/genetics.112.139337)

13. Remolina, S. C., Chang, P. L., Leips, J., Nuzhdin, S. V., Hughes, K. A. 2012 Genomic basis of aging and life-history evolution in *Drosophila melanogaster*. Evolution. 66, 3390–3403. (10.1111/j.1558-5646.2012.01710.x)

14. Carnes, M. U., Campbell, T., Huang, W., Butler, D. G., Carbone, M. A., Duncan, L. H., Harbajan, S. V., King, E. M., Peterson, K. R., Weitzel, A., et al. 2015 The Genomic Basis of Postponed Senescence in *Drosophila melanogaster*. Plos One. 10, e0138569. (10.1371/journal.pone.0138569)

15. Fabian, D. K., Garschall, K., Klepsatel, P., Santos-Matos, G., Sucena, E., Kapun, M., Lemaitre, B., Schlötterer, C., Arking, R., Flatt, T. 2018 Evolution of longevity improves immunity in Drosophila. Evol Lett. 2, 567–579. (10.1002/evl3.89)

16. Reed, L. K., Lee, K., Zhang, Z., Rashid, L., Poe, A., Hsieh, B., Deighton, N., Glassbrook, N., Bodmer, R., Gibson, G. 2014 Systems genomics of metabolic phenotypes in wild-type Drosophila melanogaster. Genetics. 197, 781–793. (10.1534/genetics.114.163857)

17. Martins, N. E., Faria, V. G., Nolte, V., Schlötterer, C., Teixeira, L., Sucena, E., Magalhaes, S. 2014 Host adaptation to viruses relies on few genes with different cross-resistance properties. Proc Natl Acad Sci U S A. 111, 5938–5943. (10.1073/pnas.1400378111)

18. Jalvingh, K. M., Chang, P. L., Nuzhdin, S. V., Wertheim, B. 2014 Genomic changes under rapid evolution: selection for parasitoid resistance. Proc Biol Sci. 281, 20132303. (10.1098/rspb.2013.2303)

19. Jha, A. R., Miles, C. M., Lippert, N. R., Brown, C. D., White, K. P., Kreitman, M. 2015 Whole genome resequencing of experimental populations reveals polygenic basis of egg size variation in Drosophila melanogaster. Mol Biol Evol. 32, 2616–2632.

20. Kang, L., Aggarwal, D. D., Rashkovetsky, E., Korol, A. B., Michalak, P. 2016 Rapid genomic changes in *Drosophila melanogaster* adapting to desiccation stress in an experimental evolution system. BMC Genomics. 17, 233. (10.1186/s12864-016-2556-y)

21. Hardy, C. M., Burke, M. K., Everett, L. J., Han, M. V., Lantz, K. M., Gibbs, A. G. 2018 Genome-Wide Analysis of Starvation-Selected *Drosophila melanogaster*-A Genetic Model of Obesity. Mol Biol Evol. 35, 50–65. (10.1093/molbev/msx254)

22. Hoedjes, K. M., van den Heuvel, J., Kapun, M., Keller, L., Flatt, T., Zwaan, B. J. 2019 Distinct genomic signals of lifespan and life history evolution in response to postponed reproduction and larval diet in *Drosophila*. Evol Lett. 3, 598–609. (10.1002/evl3.143)

23. Barrett, R. D., Hoekstra, H. E. 2011 Molecular spandrels: tests of adaptation at the genetic level. Nat Rev Genet. 12, 767–780. (10.1038/nrg3015)

24. Morris, J. Z., Navarro, C., Lehmann, R. 2003 Identification and analysis of mutations in *bob*, *Doa* and eight new genes required for oocyte specification and development in *Drosophila melanogaster*. Genetics. 164, 1435–1446.

25. Kpebe, A., Rabinow, L. 2008 Dissection of darkener of apricot kinase isoform functions in Drosophila. Genetics. 179, 1973–1987. (10.1534/genetics.108.087858)

26. Kpebe, A., Rabinow, L. 2008 Alternative promoter usage generates multiple evolutionarily conserved isoforms of *Drosophila* DOA kinase. Genesis. 46, 132–143. (10.1002/dvg.20374)

27. Rabinow, L., Samson, M.-L. 2010 The role of the *Drosophila* LAMMER protein kinase DOA in somatic sex determination. Journal of genetics. 89, 271–277.

28. Fumey, J., Wicker-Thomas, C. 2017 Mutations at the *Darkener of Apricot* locus modulate pheromone production and sex behavior in *Drosophila melanogaster*. J Insect Physiol. 98, 182–187. (10.1016/j.jinsphys.2017.01.005)

29. Yun, B., Farkas, R., Lee, K., Rabinow, L. 1994 The *Doa* locus encodes a member of a new protein kinase family and is essential for eye and embryonic development in *Drosophila melanogaster*. Genes Dev. 8, 1160–1173. (10.1101/gad.8.10.1160)

30. James, B. P., Staatz, W. D., Wilkinson, S. T., Meuillet, E., Powis, G. 2009 Superoxide dismutase is regulated by LAMMER kinase in *Drosophila* and human cells. Free Radical Biology and Medicine. 46, 821–827. (10.1016/j.freeradbiomed.2008.12.012)

31. Huang, W., Campbell, T., Carbone, M. A., Jones, W. E., Unselt, D., Anholt, R. R. H., Mackay, T. F. C. 2020 Context-dependent genetic architecture of *Drosophila* life span. PLOS Biology. 18, (10.1371/journal.pbio.3000645)

32. Ivanov, D. K., Escott-Price, V., Ziehm, M., Magwire, M. M., Mackay, T. F. C., Partridge, L., Thomton, J. M. 2015 Longevity GWAS Using the *Drosophila* Genetic Reference Panel. J Gerontol a-Biol. 70, 1470–1478. (10.1093/gerona/glv047)

33. Pallares, L. F., Lea, A. J., Han, C., Filippova, E. V., Andolfatto, P., Ayroles, J. F. 2020 Diet unmasks genetic variants that regulate lifespan in outbred *Drosophila*. bioRxiv. (10.1101/2020.10.19.346312)

34. May, C. M., van den Heuvel, J., Doroszuk, A., Hoedjes, K. M., Flatt, T., Zwaan, B. J. 2019 Adaptation to developmental diet influences the response to selection on age at reproduction in the fruit fly. J Evol Biol. 32, 425–437. (10.1111/jeb.13425)

35. Stern, D. L. 2011 Evolution, development, & the predictable genome. Greenwood Village, CO: Roberts & Co. Publishers.

36. Stern, D. L. 2000 Evolutionary developmental biology and the problem of variation. Evolution. 54, 1079–1091. (10.1111/j.0014-3820.2000.tb00544.x)

37. Hoedjes, K. M., Kostic, H., Flatt, T., Keller, L. 2021 A single nucleotide variant in the PPARgamma-homolog *Eip75B* affects fecundity in *Drosophila*. bioRxiv. (10.1101/2021.12.07.471536)

38. Osterwalder, T., Yoon, K. S., White, B. H., Keshishian, H. 2001 A conditional tissue-specific transgene expression system using inducible GAL4. Proc Natl Acad Sci U S A. 98, 12596–12601. (10.1073/pnas.221303298)

39. Tricoire, H., Battisti, V., Trannoy, S., Lasbleiz, C., Pret, A.-M., Monnier, V. 2009 The steroid hormone receptor EcR finely modulates *Drosophila* lifespan during adulthood in a sex-specific manner. Mechanisms of Ageing and Development. 130, 547–552. (10.1016/j.mad.2009.05.004)

40. Garschall, K., Dellago, H., Galikova, M., Schosserer, M., Flatt, T., Grillari, J. 2017 Ubiquitous overexpression of the DNA repair factor dPrp19 reduces DNA damage and extends *Drosophila* life span. NPJ Aging Mech Dis. 3, 5. (10.1038/s41514-017-0005-z)

41. Smith, G. D., Ebrahim, S. 2003 ’Mendelian randomization’: can genetic epidemiology contribute to understanding environmental determinants of disease? Int J Epidemiol. 32, 1–22. (10.1093/ije/dyg070)

42. Betancourt, N. J., Rajpurohit, S., Durmaz, E., Fabian, D. K., Kapun, M., Flatt, T., Schmidt, P. 2021 Allelic polymorphism at foxo contributes to local adaptation in *Drosophila melanogaster*. Mol Ecol. 30, 2817–2830. (10.1111/mec.15939)

43. Durmaz, E., Rajpurohit, S., Betancourt, N., Fabian, D. K., Kapun, M., Schmidt, P., Flatt, T. 2019 A clinal polymorphism in the insulin signaling transcription factor *foxo* contributes to life-history adaptation in *Drosophila*. Evolution. (10.1111/evo.13759)

44. Mackay, T. F. C., Richards, S., Stone, E. A., Barbadilla, A., Ayroles, J. F., Zhu, D. H., Casillas, S., Han, Y., Magwire, M. M., Cridland, J. M., et al. 2012 The *Drosophila melanogaster* Genetic Reference Panel. Nature. 482, 173–178. (10.1038/nature10811)

45. Weir, B. S., Cockerham, C. C. 1984 Estimating F-Statistics for the Analysis of Population Structure. Evolution. 38, 1358–1370. (10.1111/j.1558-5646.1984.tb05657.x)

46. Kapun, M., Nunez, J. C. B., Bogaerts-Marquez, M., Murga-Moreno, J., Paris, M., Outten, J., Coronado-Zamora, M., Tern, C., Rota-Stabelli, O., Garcia Guerreiro, M. P., et al. 2021 *Drosophila* Evolution over Space and Time (DEST) - A New Population Genomics Resource. Mol Biol Evol. (10.1093/molbev/msab259)

47. Durmaz, E., Benson, C., Kapun, M., Schmidt, P., Flatt, T. 2018 An inversion supergene in *Drosophila* underpins latitudinal clines in survival traits. J Evol Biol. 31, 1354–1364. (10.1111/jeb.13310)

48. Fabian, D. K., Kapun, M., Nolte, V., Kofler, R., Schmidt, P. S., Schlötterer, C., Flatt, T. 2012 Genome-wide patterns of latitudinal differentiation among populations of *Drosophila melanogaster* from North America. Molecular Ecology. 21, 4748–4769. (DOI 10.1111/j.1365-294X.2012.05731.x)

49. Kapun, M., Barron, M. G., Staubach, F., Obbard, D. J., Wiberg, R. A. W., Vieira, J., Goubert, C., Rota-Stabelli, O., Kankare, M., Bogaerts-Marquez, M., et al. 2020 Genomic Analysis of European *Drosophila melanogaster* Populations Reveals Longitudinal Structure, Continent-Wide Selection, and Previously Unknown DNA Viruses. Mol Biol Evol. 37, 2661–2678. (10.1093/molbev/msaa120)

50. Schmidt, P. S., Matzkin, L., Ippolito, M., Eanes, W. F. 2005 Geographic variation in diapause incidence, life-history traits, and climatic adaptation in *Drosophila melanogaster*. Evolution. 59, 1721–1732.

51. Lehtovaara, A., Schielzeth, H., Flis, I., Friberg, U. 2013 Heritability of life span is largely sex limited in *Drosophila*. Am Nat. 182, 653–665. (10.1086/673296)

52. Regan, J. C., Khericha, M., Dobson, A. J., Bolukbasi, E., Rattanavirotkul, N., Partridge, L. 2016 Sex difference in pathology of the ageing gut mediates the greater response of female lifespan to dietary restriction. eLife. 5, e10956. (10.7554/eLife.10956)

