## Supplementary Files for "Natural alleles at the *Doa* locus underpin evolutionary changes in *Drosophila* lifespan and fecundity"

***Supplementary File 1. Functional characterization of Doa*** *(with selection of references)*

| **Function/phenotype** | **Reference** |
|---|---|
| ***Functions uncovered through mutant analyses*** | |
| Oocyte formation | [1] |
| Germline differentiation in testes (through *bam*) | [2] |
| Alternative splicing (through SR proteins) | [3] |
| Sex determination, courtship behaviour, pheromones (through *transformer, doublesex*) | [3]<br>[4, 5] |
| Embryonic and post-embryotic development | [6, 7] |
| Organelle transport (through *EF1γ*) | [8] |
| Paraquat resistance (through *SOD1*) | [9] |
| Resistance to bacteria (through *Toll* pathway) | [10] |
| Cell cycle G1 phase | [11] |
| Autophagy | [12] |
| Protein secretion | [13] |
| Ecdysone signalling (through *E93*) | [6] |
| *Tor* signalling/metabolism (through *TORC1*) | [14] |
| Startle behaviour (P-element insertion screen) | [15] |
| Lifespan (P-element insertion screen, **no** effect) | [16] |
| ***Natural Doa polymorphisms associated with phenotypes (E&R and GWAS)*** | |
| Lifespan (*Doa* ranked among top50 candidate genes) | [17] |
| Aggression (confirmed with P-element insertion) | [18, 19] |
| Alcohol sensitivity (confirmed with P-element insertion) | [20] |
| Egg size, ovariole number, egg chamber length | [21] |
| Lifespan | [22] |
| Development time | [23] |
| Sleep | [24] |
| Cold acclimation | [25] |
| Lifespan GWAS (also lowered expression of *Doa* in older flies) | [26] |
| Lifespan (confirmed with RNAi) | [27] |

***Supplementary File 2. Confirmation of functioning of Doa RNAi constructs using qPCR***

Transgenic RNAi was performed using the ubiquitously expressing, mifepristone (RU486)-inducible *daughterless* (*da*)-GeneSwitch(GS)-GAL4 [28] (courtesy of Véronique Monnier, Paris) to drive the expression of Doa UAS-RNAi constructs during the adult stage. Two independent UAS-RNAi constructs obtained from the Vienna *Drosophila* RNAi Center (VDRC) were used (#19066 [D19] and #102520 [D10]). The isogenic host strain for the VDRC RNAi library, $w^{1118}$ (#60000) was used as a control to test for adverse effects of mifespristone treatment. We also tested if both RNAi lines (D10 and D19) achieved a significant knockdown of *Doa* expression using quantitative real-time PCR (qRT-PCR).

For this, we sampled female flies that had been kept on either control medium or medium with mifepristone (100 or 200 µg/ml) for four days. The flies were flash frozen in liquid nitrogen and three individuals were pooled together and homogenized for whole-body RNA extraction. RNA extractions were done using a MagMax robot, using the MagMax-96 total RNA isolation kit (ThermoFisher Scientific), following the instructions of the manufacturer. The GoScript Reverse Transcriptase system (Promega) was used to prepare cDNA; qRT-PCR was done on a QuantStudio 6 real-time PCR machine (ThermoFisher Scientific), using Power SYBR Green PCR Master mix (ThermoFisher Scientific). We analyzed ten biological samples per construct and mifepristone treatment, with three technical qRT-PCR replicates each. The expression levels of *Doa* and four reference genes (*TBP*, *Cyp1*, *Ef1a48D* and *Rap2l*; see Table 1 for information on the primers used) were measured. The reference genes had a stability of M<0.25 as determined from a set of female whole-body samples of various strains and mifepristone concentrations using the GeNorm algorithm incorporated in qbasePLUS v.1.5 (Biogazelle). The gene expression levels of the four reference genes were averaged geometrically to obtain an accurate normalization factor for each sample (Vandesompele et al, 2002). The relative gene expression levels of *Doa* for the two RNAi constructs were then calculated compared to the expression of the control treatment (i.e. no mifepristone) for both constructs, respectively (Table 2, Figure 1). The gene expression levels were analyzed in R (v.3.3.1) using linear mixed models (package *lme4*, v.1.1-13) with mifepristone concentration as a fixed effect (continuous variable) and batch (two sets of five samples were measured in parallel) as random factor.

We detected a significant reduction in whole-body *Doa* expression by both RNAi constructs when being fed mifepristone (D10: $\chi 2$ = 7.51, *P* = 0.0061; D19: $\chi 2$ = 13.06, *P* = 0.0003). For the construct D10, there was an average reduction in *Doa* expression of 13% at a mifepristone concentration of 200 µg/ml. For D19 this reduction was 17%.

**Table 1:** primers used for qRT-PCR.

| Gene | Forward primer | Reverse primer | Amplicon (bp) | Primer efficiency |
|---|---|---|---|---|
| *Doa* | CCGTGATCACTGCAAGCCGTTGT | TGTGGTGGCAGCCTATCAAAGAACG | 160 | 1.92 |
| *TBP* | CGGTTTCCCTGCAAAGTTCCTCGA | CACGATTCGAGGTCGCACCATACG | 172 | 1.92 |
| *Cyp1* | GTCGGCAGCGGCATTTCAGAT | CTGCACGCTGACGAAGCTAGG | 78 | 1.91 |
| *Ef1a48D* | TGCCACACCGCTCACATTGCT | CACGCACAGGGGCTTAGAGG | 150 | 1.96 |
| *Rap2l* | TATTGGACACCGCGGGCACA | TGGGTGCTGGCTGACTTCCT | 168 | 1.96 |

**qPCR protocol: 95°C for 10 min; 40 cycles with 95°C for 15 sec and 60°C for 1 min; melting analysis from 60°C until 95°C (+0.05 °C/s).*

**Table 2:** Changes in *Doa* relative gene expression after mifepristone treatment (compared to control, i.e. 0 RU).

| Acronym | Construct | Relative expression | | $\chi 2$ | *P* |
|---|---|---|---|---|---|
| | | 100 RU | 200 RU | | |
| D10 | VDRC #102520 | 0.88 | 0.87 | 7.51 | 0.0061 |
| D19 | VCDR #19066 | 0.87 | 0.83 | 13.06 | 0.00030 |

**Figure 1:** Changes in *Doa* gene expression after mifepristone treatment. The relative expression of *Doa* of the two mifepristone concentrations compared the control (no mifepristone) is shown.

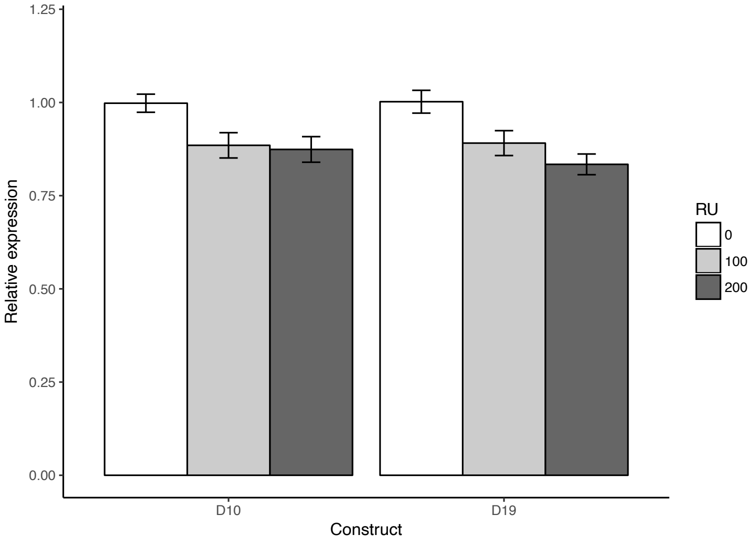

***Supplementary File 3. Effects of mifepristone exposure on lifespan and reproduction of da-GS-GAL4/w$^{1118}$ controls***

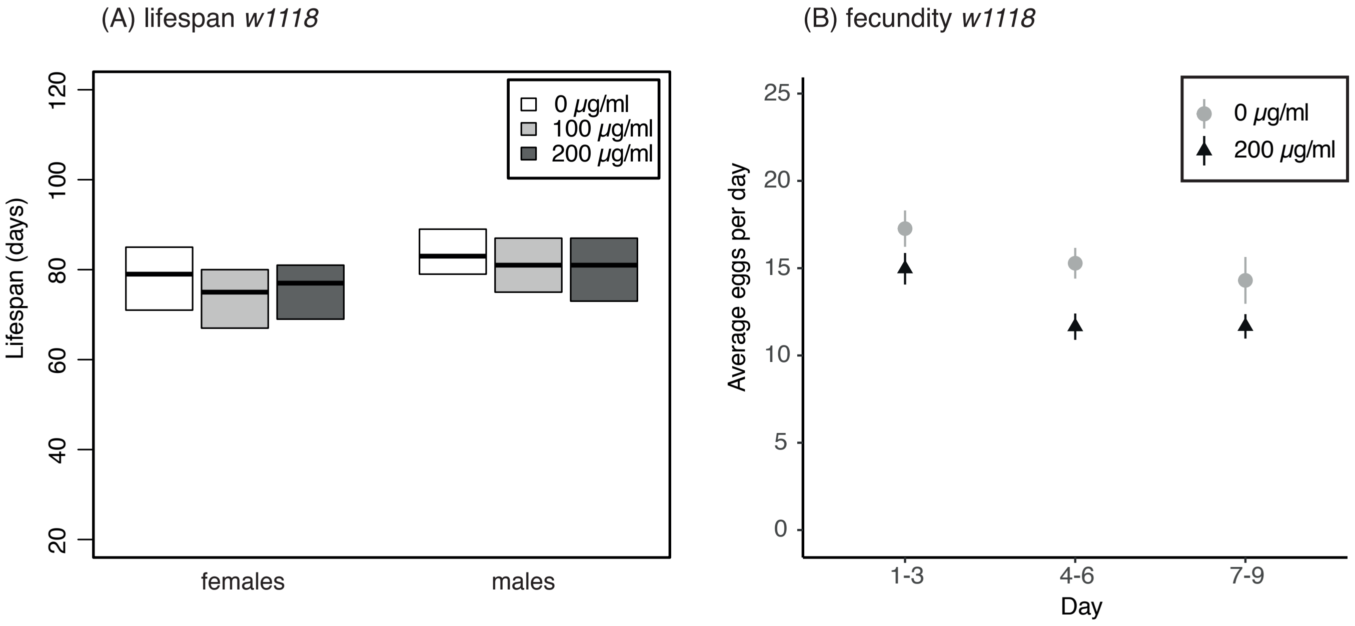

**Figure 1:** Effects of mifepristone exposure (0, 100, or 200 µg/ml) on lifespan and fecundity of the control strain.

#### ***Supplementary File 4. Linkage disequilibrium among Doa SNPs in the DGRP***

To estimate potential confounding effects of other SNPs, located closely to our four *Doa* candidate SNPs, in our Mendelian randomization approach, we estimated LD (measured by pairwise $r^2$) as done by [29]) for all SNPs (minor allele frequency ≥ 0.1) located within *Doa* in the complete panel of DGRP lines. These analyses indicate very low levels of LD across the entire gene (Figure 1), but also among the four candidate SNPs (Table 1). The Mendelian randomization approach is, therefore, expected to provide information on the functional impact of a candidate SNP, with little or no confounding effects of other (*Doa*) SNPs.

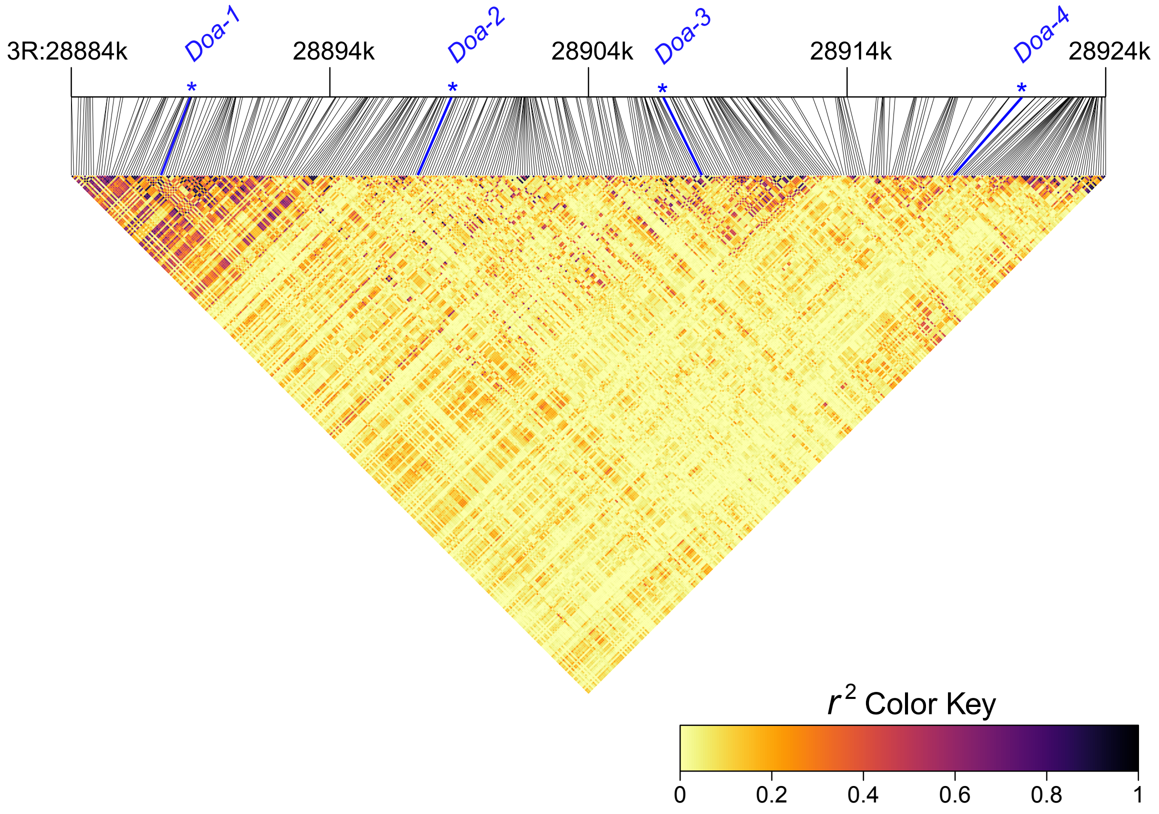

**Figure 1:** Linkage disequilibrium (measured by pairwise $r^2$) of SNPs within *Doa* in the DGRP. Coordinates according to genome reference v6.

**Table 1:** Pairwise linkage disequilibrium (measured by pairwise $r^2$) among the four candidate *Doa* SNPs in the DGRP.

| | *Doa-1* | *Doa-2* | *Doa-3* | *Doa-4* |
|---|---|---|---|---|
| *Doa-1* | x | 0.234 | 0.027 | 0.169 |
| *Doa-2* | | x | 0.095 | 0.121 |
| *Doa-3* | | | x | 0.029 |
| *Doa-4* | | | | x |

### *Supplementary File 5. SNP-wise differentiation among panels with alternative allelic state of the SNP association study*

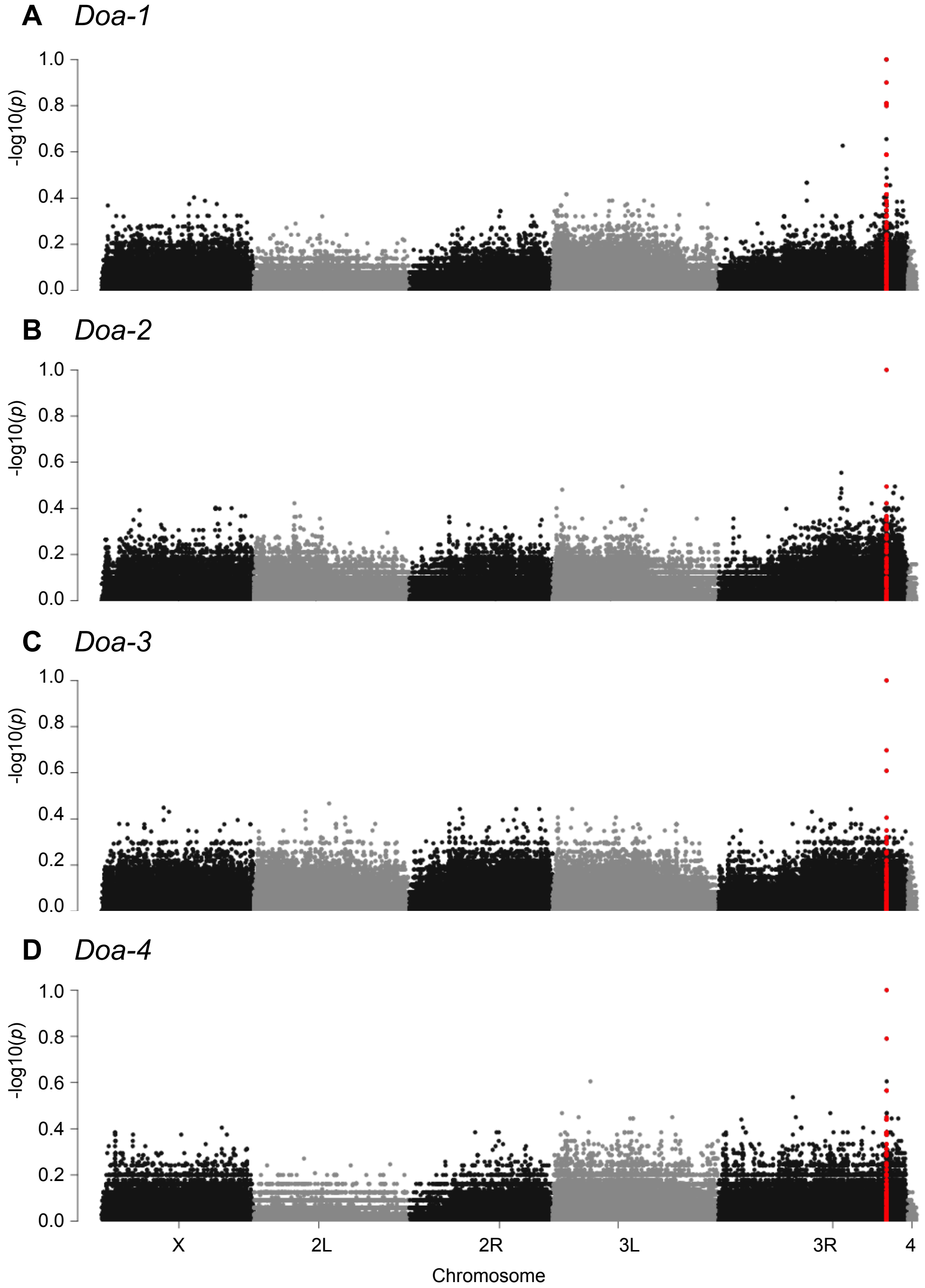

**Figure 1:** Genetic differentiation ($F_{ST}$) between two panels with alternative allelic state was calculated for each of the four *Doa* SNPs by pooling genome sequence information of all lines per panel. These analyses indicate that only the candidate SNP is fixed ($F_{ST}$ = 1) between the two panels, and that very few SNPs show a high differentiation ($F_{ST}$ > 0.5) between the two panels. This is observed for each of the four *Doa* SNPs. This indicates that there were little or no confounding genetic factors in our SNP association study.

***Supplementary File 6. Population genetic analysis of Doa SNP alleles***

*Methodology of the population genetic analysis*

We examined the allele frequencies of the four candidate *Doa* SNPs in natural populations from Europe, North-America, and Africa. For this, we analyzed whole genome sequencing data from multiple geographic populations available from the DEST dataset [30]. In short, this dataset consisted of single individual sequencing data and Pool-Seq data from various sources that was re-mapped and quality filtered in a synchronized fashion. For the single individual sequencing data, the allele frequency of the *Doa* alleles was calculated when at least ten individuals were sequenced for a given population. Data from pooled populations was included when the coverage for each *Doa* SNP was at least 10. This data provides an overview of the distribution of each pair of alleles worldwide. In addition, we analyzed the PoolSeq data from the 172 samples collected in Europe to test the clinal distribution of the allele frequencies. For this, we used generalized linear models (GLMs) based on the allele counts of the *Doa* SNPs, with either longitude or latitude as explanatory factor. To estimate false discovery rate, we also tested clinality of a set of 21'008 neutral SNPs [31, 32]. Empirical cumulative density functions (ECDF) were generated based on the $\chi 2$ values of these neutral SNPs. The area of the upper tail confined by the $\chi 2$ values of the *Doa* SNPs indicates the percentile of neutral SNPs with $\chi 2$ values equal or higher than the *Doa* SNPs. This parameter provides a corrected estimate of significance of clinality of each SNP compared to genome-wide neutral estimates [33].

*Distribution of Doa SNP alleles*

The allelic variants of each of the four *Doa* SNPs identified in the experimental evolution study by [34] were also observed in natural populations across Europe and North America, with both alleles of each SNP co-occurring at all geographical locations sampled (Allele frequencies observed: *Doa-1*: 0.16-0.57, *Doa-2*: 0.34-0.72, *Doa-3*: 0.65-0.98, *Doa-4*: 0.51-0.95) (Figure 2, Supplementary Table 2). This confirms that these four SNP positions are polymorphic in natural populations as well, and may therefore be targets of natural selection in these populations.

A significant clinal pattern in the allele frequency distribution among European populations was observed for two of the SNP positions. The allele frequencies of

*Doa-3* are distributed longitudinally with on average higher frequencies of the "G" allele, associated with postponed reproduction and longer lifespan in the EE study, in eastern European populations. This observation corresponds with the longitudinal population structure observed in European *D. melanogaster*, and may indicate a role in local climate adaptation [35]. The "G" allele is pre-dominant in the African populations studied (AF: 0.9-1), which may indicate that it is the ancestral allele.

For *Doa-4* we observed a significant latitudinal distribution with a higher frequency of the "T" allele, associated with postponed reproduction and longevity in the EE study, in northern European populations. The “T” allele frequencies of *Doa-4* ranged between 0.51 and 0.95 in the European populations sampled, whereas the "T" variant was the minor allele in the African populations (AF: 0-0.4), which may support the idea that this allele provides an adaptation to climatic conditions at higher latitudes, which includes adaptation of life history [29, 36].

**Table 1:** Statistical analyses of the latitudinal and longitudinal distribution of *Doa* alleles.

| | Latitude | | | Longitude | | |
|---|---|---|---|---|---|---|
| | $\chi 2$ | *P* | corrected *P* | $\chi 2$ | *P* | corrected *P* |
| *Europe* | | | | | | |
| *Doa-1* | 13.6 | 2.23E-04 | 0.25 | 32.7 | 1.09E-08 | 0.27 |
| *Doa-2* | 17.9 | 2.32E-05 | 0.21 | 53.9 | 2.07E-13 | 0.20 |
| *Doa-3* | 31.7 | 1.84E-08 | 0.14 | 222.9 | <2.2E-16 | **0.037** |
| *Doa-4* | 174.3 | 2.2E-16 | **0.0069** | 43.7 | 3.91E-11 | 0.23 |

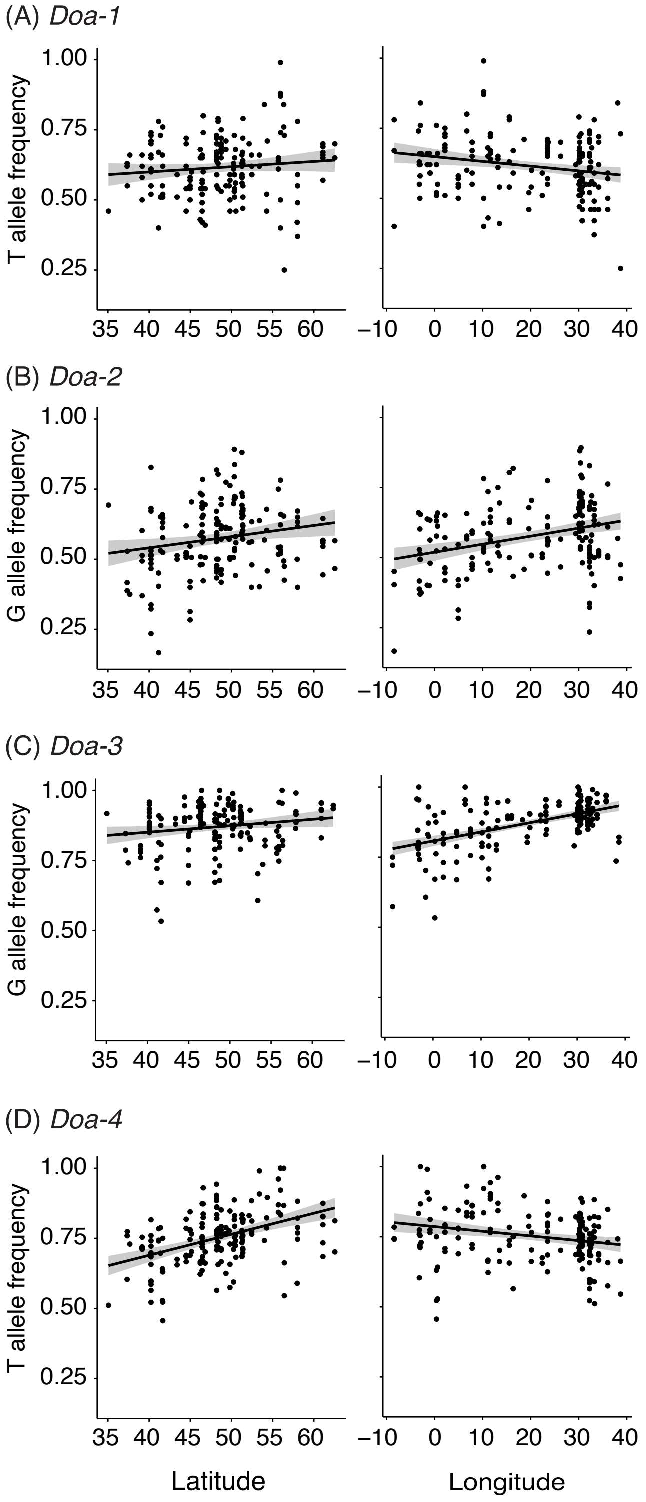

**Figure 1:** Latitudinal and longitudinal distribution of the allele frequencies of *Doa-1*, *Doa-2*, *Doa-3*, and *Doa-4* among European populations.

### *References*

1. Morris, J.Z., C. Navarro, and R. Lehmann, *Identification and analysis of mutations in bob, Doa and eight new genes required for oocyte specification and development in Drosophila melanogaster.* Genetics, 2003. **164**(4): p. 1435-46.
2. Zhao, S.W., et al., *The highly conserved LAMMER/CLK2 protein kinases prevent germ cell overproliferation in Drosophila.* Developmental Biology, 2013. **376**(2): p. 163-170.
3. Kpebe, A. and L. Rabinow, *Dissection of darkener of apricot kinase isoform functions in Drosophila.* Genetics, 2008. **179**(4): p. 1973-1987.
4. Fumey, J. and C. Wicker-Thomas, *Mutations at the Darkener of Apricot locus modulate pheromone production and sex behavior in Drosophila melanogaster.* J Insect Physiol, 2017. **98**: p. 182-187.
5. Rabinow, L. and M.-L. Samson, *The role of the Drosophila LAMMER protein kinase DOA in somatic sex determination.* J Genet, 2010. **89**: p. 271-277.
6. Kpebe, A. and L. Rabinow, *Alternative promoter usage generates multiple evolutionarily conserved isoforms of Drosophila DOA kinase.* Genesis, 2008. **46**(3): p. 132-143.
7. Yun, B., et al., *The Doa locus encodes a member of a new protein kinase family and is essential for eye and embryonic development in Drosophila melanogaster.* Genes Dev, 1994. **8**(10): p. 1160-73.
8. Serpinskaya, A.S., et al., *Protein kinase Darkener of apricot and its substrate EF1gamma regulate organelle transport along microtubules.* J Cell Sci, 2014. **127**(Pt 1): p. 33-9.
9. James, B.P., et al., *Superoxide dismutase is regulated by LAMMER kinase in Drosophila and human cells.* Free Radical Biology and Medicine, 2009. **46**(6): p. 821-827.
10. Kanoh, H., et al., *Ex vivo genome-wide RNAi screening of the Drosophila Toll signaling pathway elicited by a larva-derived tissue extract.* Biochem Biophys Res Commun, 2015. **467**(2): p. 400-6.
11. Bettencourt-Dias, M., et al., *Genome-wide survey of protein kinases required for cell cycle progression.* Nature, 2004. **432**(7020): p. 980-7.
12. Gorski, S.M., et al., *A SAGE approach to discovery of genes involved in autophagic cell death.* Curr Biol, 2003. **13**(4): p. 358-63.
13. Bard, F., et al., *Functional genomics reveals genes involved in protein secretion and Golgi organization.* Nature, 2006. **439**(7076): p. 604-7.
14. Tang, H.W., et al., *The TORC1-Regulated CPA Complex Rewires an RNA Processing Network to Drive Autophagy and Metabolic Reprogramming.* Cell Metab, 2018. **27**(5): p. 1040-1054 e8.
15. Yamamoto, A., et al., *Neurogenetic networks for startle-induced locomotion in Drosophila melanogaster.* Proc Natl Acad Sci U S A, 2008. **105**(34): p. 12393-8.
16. Magwire, M.M., et al., *Quantitative and molecular genetic analyses of mutations increasing Drosophila life span.* PLoS Genet, 2010. **6**(7): p. e1001037.
17. Ivanov, D.K., et al., *Longevity GWAS Using the Drosophila Genetic Reference Panel.* Journals of Gerontology Series a-Biological Sciences and Medical Sciences, 2015. **70**(12): p. 1470-1478.
18. Edwards, A.C., et al., *Quantitative genomics of aggressive behavior in Drosophila melanogaster.* PLoS Genet, 2006. **2**(9): p. e154.

19. Zwarts, L., et al., *Complex genetic architecture of Drosophila aggressive behavior.* Proceedings of the National Academy of Sciences of the United States of America, 2011. **108**(41): p. 17070-17075.
20. Morozova, T.V., T.F.C. Mackay, and R.R.H. Anholt, *Transcriptional Networks for Alcohol Sensitivity in Drosophila melanogaster.* Genetics, 2011. **187**(4): p. 1193-U345.
21. Jha, A.R., et al., *Whole genome resequencing of experimental populations reveals polygenic basis of egg size variation in Drosophila melanogaster.* Mol Biol Evol, 2015. **32**(10): p. 2616-2632.
22. Fabian, D.K., et al., *Evolution of longevity improves immunity in Drosophila.* Evol Lett, 2018. **2**(6): p. 567-579.
23. Burke, M.K., et al., *Genome-wide analysis of a long-term evolution experiment with Drosophila.* Nature, 2010. **467**(7315): p. 587-U111.
24. Svetec, N., et al., *Evidence that natural selection maintains genetic variation for sleep in Drosophila melanogaster.* BMC Evol Biol, 2015. **15**: p. 41.
25. MacMillan, H.A., et al., *Cold acclimation wholly reorganizes the Drosophila melanogaster transcriptome and metabolome.* Sci Rep, 2016. **6**: p. 28999.
26. Pallares, L.F., et al., *Diet unmasks genetic variants that regulate lifespan in outbred Drosophila.* bioRxiv, 2020.
27. Huang, W., et al., *Context-dependent genetic architecture of Drosophila life span.* PLOS Biology, 2020. **18**(3).
28. Tricoire, H., et al., *The steroid hormone receptor EcR finely modulates Drosophila lifespan during adulthood in a sex-specific manner.* Mechanisms of Ageing and Development, 2009. **130**(8): p. 547-552.
29. Durmaz, E., et al., *A clinal polymorphism in the insulin signaling transcription factor foxo contributes to life-history adaptation in Drosophila.* Evolution, 2019.
30. Kapun, M., et al., *Drosophila Evolution over Space and Time (DEST) - A New Population Genomics Resource.* Mol Biol Evol, 2021.
31. Parsch, J., et al., *On the utility of short intron sequences as a reference for the detection of positive and negative selection in Drosophila.* Mol Biol Evol, 2010. **27**(6): p. 1226-34.
32. Clemente, F. and C. Vogl, *Unconstrained evolution in short introns? - an analysis of genome-wide polymorphism and divergence data from Drosophila.* J Evol Biol, 2012. **25**(10): p. 1975-1990.
33. Ramaekers, A., et al., *Altering the Temporal Regulation of One Transcription Factor Drives Evolutionary Trade-Offs between Head Sensory Organs.* Dev Cell, 2019. **50**(6): p. 780-792 e7.
34. Hoedjes, K.M., et al., *Distinct genomic signals of lifespan and life history evolution in response to postponed reproduction and larval diet in Drosophila.* Evol Lett, 2019.
35. Kapun, M., et al., *Genomic Analysis of European Drosophila melanogaster Populations Reveals Longitudinal Structure, Continent-Wide Selection, and Previously Unknown DNA Viruses.* Mol Biol Evol, 2020. **37**(9): p. 2661-2678.
36. Fabian, D.K., et al., *Genome-wide patterns of latitudinal differentiation among populations of Drosophila melanogaster from North America.* Molecular Ecology, 2012. **21**(19): p. 4748-4769.
